# Head-to-head comparison of four antigen-based rapid detection tests for the diagnosis of SARS-CoV-2 in respiratory samples

**DOI:** 10.1101/2020.05.27.119255

**Authors:** Thomas Weitzel, Paulette Legarraga, Mirentxu Iruretagoyena, Gabriel Pizarro, Valeska Vollrath, Rafael Araos, José M. Munita, Lorena Porte

## Abstract

In the context of the Covid-19 pandemic, the development and validation of rapid and easy-to-perform diagnostic methods are of high priority. We compared the performance of four rapid antigen detection tests for SARS-CoV-2 in respiratory samples. Immunochromatographic SARS-CoV-2 assays from RapiGEN, Liming bio, Savant, and Bioeasy were evaluated using universal transport medium containing naso-oropharyngeal swabs from suspected Covid-19 cases. The diagnostic accuracy was determined in comparison to SARS-CoV-2 RT-PCR. A total of 111 samples were included; 80 were RT-PCR positive. Median patients’ age was 40 years, 55% were female, and 88% presented within the first week after symptom onset. The evaluation of the Liming bio assay was discontinued due to insufficient performance. The overall sensitivity values of RapiGEN, Liming bio, and Bioeasy tests were 62.0% (CI95% 51.0–71.9), 16.7% (CI95% 10.0–26.5), and 85.0% (CI95% 75.6–91.2), respectively, with specificities of 100%. Sensitivity was significantly higher in samples with high viral loads (RapiGEN, 84.9%; Bioeasy, 100%). The study highlighted the significant heterogeneity of test performance among evaluated assays, which might have been influenced by the use of a non-validated sample material. The high sensitivity of some tests demonstrated that rapid antigen detection has the potential to serve as an alternative diagnostic method, especially in patients presenting with high viral loads in early phases of infection. This is particularly important in situations with limited access to RT-PCR or prolonged turnaround time. Further comparative evaluations are necessary to select products with high performance among the growing market of diagnostic tests for SARS-CoV-2.

## Introduction

Since its emergence in 2019, the SARS-CoV-2 pandemic has caused tremendous public health challenges worldwide (1). Early detection of cases is essential to help curtail this unprecedented pandemic; thus, rapid and easy-to-perform diagnostic tools that can be used to test large numbers of samples in a short period of time are crucial (2). To date, the recommended diagnostic method for SARS-CoV-2 infection (known as Covid-19) is real-time reverse-transcription polymerase chain reaction (RT-PCR), which was introduced in January 2020 (3), and is now performed using WHO or CDC protocols (4, 5), as well as various commercial assays (6).

The gap between the number of samples and the capacity to perform RT-PCR in a timely manner is considered a mayor limitation of public health containment strategies (7). Therefore, there is a critical demand for alternative assays, especially rapid diagnostic tests (RDTs), which are timely, easy to perform, and can serve for point-of-care testing (POCT) or community-based testing (8). RDTs for antibody detection have been developed, but due to the delay in humoral immune response, they have limited use for early diagnosis and low sensitivity for community-based screening (9, 10). Antigen detection tests, on the other hand, have the advantage of detecting the presence of the virus itself and might therefore be a better tool for early cases, but require sufficient viral loads and high-quality sampling (11). Although SARS-CoV-2-specific antigen testing has only recently been developed (12); the market pressure generated by this unprecedented pandemic has resulted in many novel assays that are now commercially available (13). Unfortunately, scientific literature supporting their accuracy is scarce and real-world performance of these assays is uncertain; their validation and comparison are therefore of high priority (12, 14). Here we present a head-to-head comparison and evaluation of four novel antigen-based RDTs for the detection of SARS-CoV-2 in respiratory specimens from suspected Covid-19 cases.

## Material and Methods

We conducted a head-to-head study of the diagnostic accuracy of four rapid SARS-CoV-2 antigen detection tests compared to RT-PCR. Samples derived from patients with respiratory symptoms and/or fever, attending Clínica Alemana, a private medical center in Santiago, Chile,^10^ between March 16 and April 26, 2020. Specimens were obtained by trained personnel in a specially dedicated “Respiratory Emergency Room” and consisted of naso-oropharyngeal (NOP) swabs, which were placed in tubes with 3 mL universal transport medium (UTM-RT^®^ System, Copan Diagnostics, Murrieta, CA, USA). Samples were initially examined for SARS-CoV-2 RNA by COVID-19 Genesig^®^ Real-Time PCR assay (Primerdesign Ltd., Chander’s Ford, UK) after RNA extraction with the Magna Pure Compact system (Roche Molecular Systems Inc., Pleasanton, Ca, USA) or using a manual protocol with the High Pure Viral Nucleic Acid kit (Roche Molecular Systems Inc., Mannheim, Germany). The Primerdesign RT-PCR received FDA Emergency Use Authorization (EUA) and is within the WHO Emergency Use Listing (EUL) tests eligible for procurement. The test kit includes a positive control template of the target gene and a RNA internal extraction control. It targets the RNA-dependent RNA polymerase (RdRp) with a detection limit of 0.58 copies/μL, according to the manufacturer. Samples showing an exponential amplification curve and a cycle threshold (Ct) value ≤40 were considered as positive.

PCR-characterized samples (UTM with swabs) were kept at −80° C and tested on April 28 and 29 by the following lateral flow antigen-detection kits: 1) “Biocredit COVID-19 Ag One Step SARS-CoV-2 Antigen Test” (RapiGEN Inc., Anyang-si, Gyeonggi-do, Republic of Korea), 2) “COVID-19 Antigen Rapid Test Device StrongStep® COVID-19 Antigen Test” (Liming Bio-Products Co., Jiangsu, China; 3) “Huaketai New Coronavirus (SARS-CoV-2) N Protein Detection Kit (Fluorescence immunochromatography)” (Savant Biotechnology Co., Beijing, China), and 4) “Diagnostic Kit for 2019-Novel Coronavirus (2019-nCoV) Ag Test (Fluorescence Immunochromatographic Assay)” (Bioeasy Biotechnology Co., Shenzhen, China). All kits have a cassette format and display test and control lines, permitting a rapid use without positive and negative control specimens (Table 1). Tests must be read after a specific incubation period (5 to 15 minutes). The first two assays use colloid gold conjugated antibodies, resulting in visible colored bands, whereas the latter two kits are based on fluorescein-marked antibodies. For the Savant assay we used a UV flashlight provided by the manufacturer for visual readout, while the Bioeasy kit was automatically read by the immunofluorescence analyzer EASY-11 (Bioeasy Biotechnology Co.).

**Table 1.** Main characteristics of four rapid SARS-CoV-2 antigen-detection tests (as recommended by manufacturer)

| Characteristic | Test N°1 | Test N°2 | Test N°3 | Test N°4 |
| --- | --- | --- | --- | --- |
| Manufacturer | RapiGen | Liming bio | Savant | Bioeasy |
| Commercial name | Biocredit One Step SARS-CoV-2 Antigen Test | StrongStep® COVID-19 Antigen Test | Huaketai New Coronavirus (SARS-CoV-2) N Protein Detection Kit (FIA) | Diagnostic Kit for 2019-Novel Coronavirus (2019-nCoV) Ag Test (FIA) |
| Catalogue N° | G61RHA20 | 500200 | BCT-HKT-050 | YRLF04401025 |
| Lot N° | H073001SD | 2003014 | 20031501 | 2002N408 |
| Certification<sup>2</sup> | CE-IVD | CE-IVD | CE-IVD | CE-IVD |
| Antigen detected | Not specified | Not specified | N protein | Not specified<sup>1</sup> |
| Specimen | NP/OP swab | NP/OP swab | Throat swab | NP/OP swab, sputum |
| Extraction/<br>Dilution | Swab immersed in assay diluent and strongly squeezed | Swab immersed in extraction buffer and strongly squeezed | Swab immersed in conservation solution immediately before testing | Swab immersed in extraction buffer and strongly squeezed |
| Volume applied<br>into cassette | 3-4 drops<br>(~90-150 µL) | 3 drops<br>(~100 µL) | 60 µL | 2 drops<br>(100 µL) |
| Study sample | 150 µL of UTM | 150 µL of UTM | 60 µL of UTM | 100 µL of UTM |
| Incubation (at<br>room temperature) | 5-8 minutes | 15-20 minutes | 15 minutes ±1 minute | 10 minutes ±0 minutes |
| Readout | Visual: colored bands | Visual: colored bands | Visual: fluorescent bands<br>(under UV light) | Automated: fluorescence<br>reader |

FIA, fluorescence immune assay; NP, nasopharyngeal; OP, oropharyngeal
<sup>1</sup> N protein (24)
<sup>2</sup> Information from (13)

Importantly, our evaluation protocol included a deviation from the manufacturer’s instructions. Instead of using the provided test solution, we used the equivalent volume of UTM, as described in other studies (15–17). This method allowed us to compare all four assays using the same specimens and to rapidly evaluate a large number of samples previously characterized by RT-PCR.

Positive and negative samples were selected by convenience among the 5,276 respiratory specimens processed for SARS-CoV-2 in the clinical laboratory during the study period. Due to the shortage of available test kits, a 2:1 distribution of positive to negative samples was chosen. Seventeen of the positive specimens had been used in a previous evaluation of the Bioeasy RDT by our group (15). Assays were tested in parallel from the same sample and by the same trained technician, who was blinded to the RT-PCR results. All test procedures, except the reading of the cassette, were performed under a BSL2 cabinet. Assays with visual output were read by two independent observers, who conferred with a third observer in case of disagreement. Results of the RDTs were compared to those of RT-PCR as reference method; for samples with discordant result, tests were repeated. Demographic and clinical data were obtained from the mandatory Covid-19 notification forms and analyzed in an anonymized manner. Samples with high viral loads (defined as Ct value <25) were compared to those with low viral load (Ct values >25) (16). Statistical analyses were performed with OpenEpi (version 3.01) and GraphPad Prism (version 8.4.2).

All materials and personnel for this evaluation except the test from Savant Biotechnology Co. were purchased using funds for routine diagnostics of the Clinical Laboratory of Clínica Alemana; the Savant RDT was provided free-of-charge by the manufacturer. The study was approved by the respective institutional review board (Comité Etico Científico, Facultad de Medicina Clínica Alemana, Universidad del Desarrollo, Santiago, Chile) and need for informed consent was waived.

## Results

The study included a total of 111 samples from symptomatic patients; 55% were female, with a median age of 40 years, which was similar to the population of all patients tested for Covid-19 in our laboratory during the study period (57% female, median age 38 years). Eighty of the tested specimens were RT-PCR positive for SARS-CoV-2, which represented 22% of all positives during the study period, while 31 samples were RT-PCR negative. The median duration from symptom onset to sampling was 2 days (IQR 1-5 days); 88% of specimens (96/109; missing data, n=2) were taken during the first week of symptoms. Ct values ranged from 10.7 to 37.7 (mean, 22.5).

While all four tests were user-friendly, test performances showed significant differences (Table 2). The evaluation of the Liming bio kit was stopped after 19 samples, due to its poor results with a sensitivity of 0% (0/9), specificity of 90% (9/10), and a Kappa coefficient of −0.1. Sensitivities of the other three assays ranged from 16.7% for the Savant test to 85% for Bioeasy (Table 2). Similarly, Bioeasy had the highest accuracy (89.2%) and Kappa coefficient (0.8). All three assays had a specificity of 100% and proved robust; invalid results occurred only with RapiGen (n=2) and Savant (n=2) due to insufficient liquid migration.

**Table 2.** Performance of four antigen detection tests for SARS-CoV-2 compared to RT-PCR

| Antigen detection test |  | RT-PCR |  | Sensitivity |  | Specificity |  | Accuracy | Kappa |
| --- | --- | --- | --- | --- | --- | --- | --- | --- | --- |
| Assay | Result | Positive | Negative | % | CI95% | % | CI95% | % | coefficient |
| RapiGen<br>(n = 109)<sup>1</sup> | Positive | 49 | 0 | 62.0 | 51.0-71.9 | 100 | 88.7-100 | 72.5 | 0.5 |
|  | Negative | 30 | 30 |  |  |  |  |  |  |
| Liming bio<br>(n = 19)<sup>2</sup> | Positive | 0 | 1 | 0 | 0.0-29.9 | 90.0 | 59.6-98.2 | 47.4 | -0.1 |
|  | Negative | 9 | 9 |  |  |  |  |  |  |
| Savant<br>(n = 109)<sup>1</sup> | Positive | 13 | 0 | 16.7 | 10.0-26.5 | 100 | 89.0-100 | 40.4 | 0.1 |
|  | Negative | 65 | 31 |  |  |  |  |  |  |
| Bioeasy<br>(n = 111) | Positive | 68 | 0 | 85.0 | 75.6-91.2 | 100 | 89.0-100 | 89.2 | 0.8 |
|  | Negative | 12 | 31 |  |  |  |  |  |  |

<sup>1</sup> Two invalid results were excluded
<sup>2</sup> Testing was suspended after 19 samples due to poor test performance

In addition, we correlated the viral loads of samples with true positive and false negative results. As shown in Figure 1, mean Ct values of false negative samples were significantly higher than in true positives samples in all three tests. RapiGen and Bioeasy mainly missed to detect samples with high viral load (Ct values >25); this threshold was less clear for the Savant assay. Accordingly, sensitivity values of RapiGen and Bioeasy differed significantly among subgroups with high viral loads and low viral loads (Ct >25) (Figure 2). Indeed, the two assays identified 84.9% and 100% of specimens with high viral load, but only 15.4% and 53.8% of those with low viral load, respectively. In case of the Savant assay, although the difference in sensitivity among samples with high and low viral loads remained, it was far less striking (21.2% vs. 7.7%, respectively) (Figure 2). RapiGen and Bioeasy had a concordance of 82%, while agreement between these two tests and Savant was 67% and 50%, respectively. As visualized in Figure 3, all samples identified as positive by RapiGen or Savant were also detected by Bioeasy. Moreover, the latter platform identified various additional positive samples, especially among those with lower viral loads (Ct 22-30) and it was the only test to detect specimens with very low viral loads (Ct >30).

**Figure 1.**
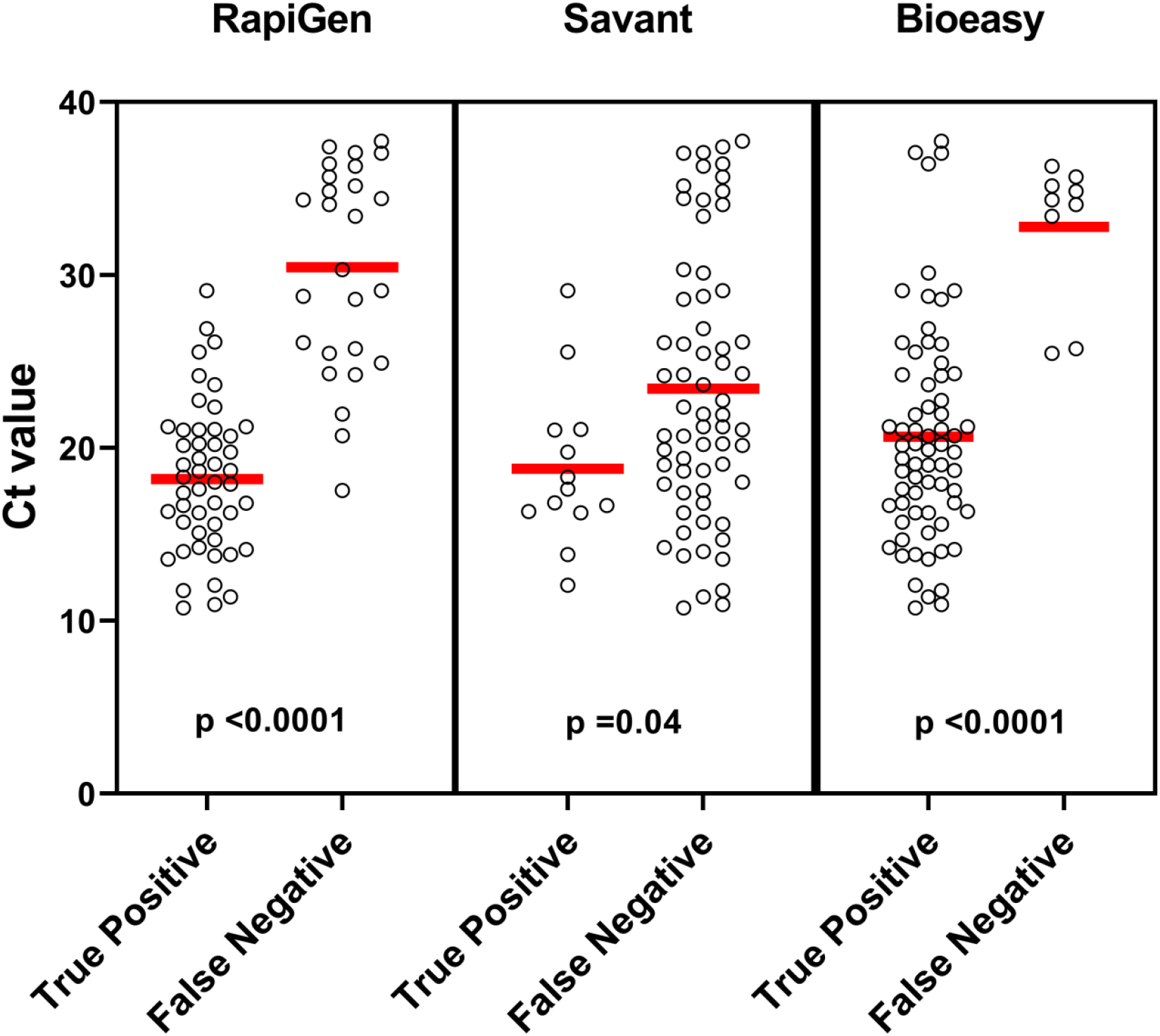
Comparison of viral loads (Ct values) among true positive and false negative results of three rapid antigen assays. Red lines represent mean values; p values calculated by two-tailed t-Test.

**Figure 2.**
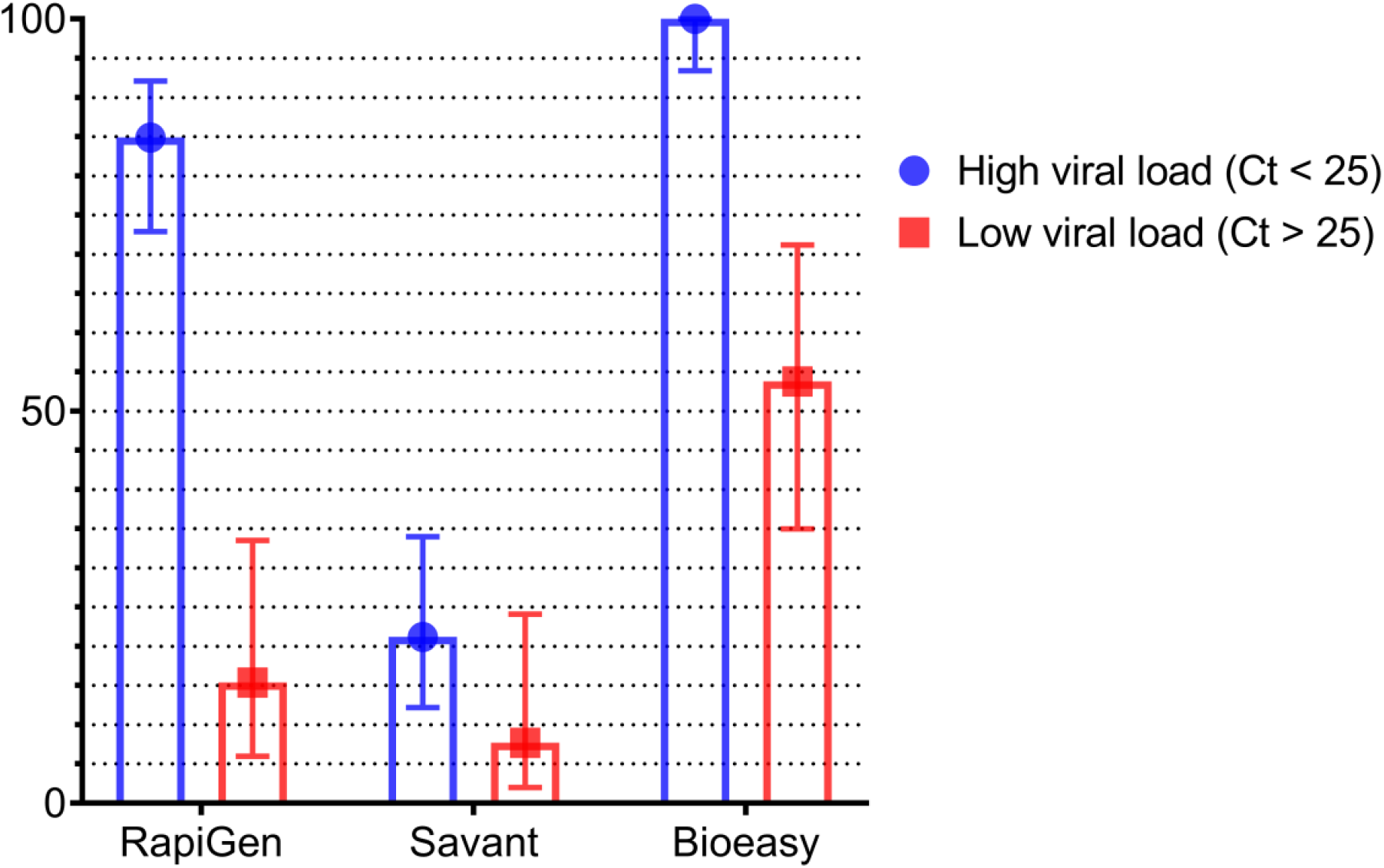
Sensitivity (%) of three rapid antigen tests in subgroups of samples with high viral load (Ct <25) and low viral load (Ct >25). Sensitivity values are represented by dots and squares, while bars demonstrate 95% confidence intervals.

**Figure 3.**
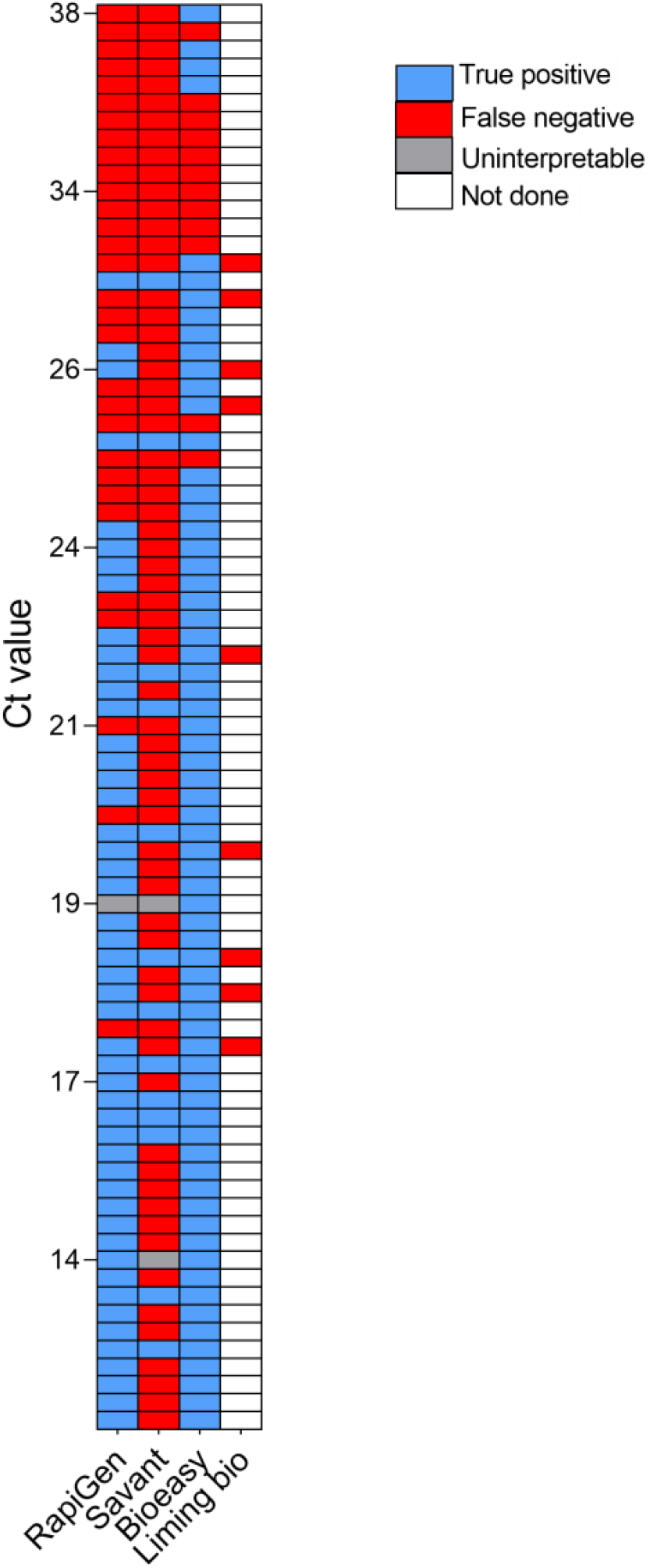
Concordance of four rapid antigen tests among 80 RT-PCR positive samples. Each line represents an individual sample. Samples are listed from high Ct values (top) to low Ct values (bottom).

All four tests were incubated on a flat surface at room temperature and delivered results in short time (5 to 15 minutes). The visual readout of RapiGen was clear, regardless of the intensity of the band, due to its dual colored format in which the control band was red and positive results appeared as black lines. The visual interpretation of the fluorescence Savant assay was difficult under daylight conditions. This test might perform differently with its reader, which was not available in Chile. Bioeasy had a user-friendly readout performed by a desktop instrument that interpreted the cassettes in <1 minute. The instrument included options for saving patient results, QR coding, and printing, and is suitable to be connected to a laboratory information system.

## Discussion

The evaluated assays are among the 22 antigen detection tests for SARS-CoV-2 available at the time of writing, of which 14 are commercialized with CE (Conformité Européene) marking (13). However, it is important to note that the CE licencing process is based on self-reporting of manufacturers, does not grant high performance, and can be misused (18, 19). Indeed, the challenge and problems presented by this procedure has recently been addressed by the European Commission in light of the evolving Covid-19 pandemic (12). The emerging marketing and use of novel PCR-independent tests in the absence of robust performance data has been criticized and it has the potential to cause more damage than benefit (20, 21). Independent evaluations of such diagnostic assays are therefore urgently needed, with larger comparative studies for antibody tests only recently available or under way (22, 23). Antigen-based testing is still in its infancy and evaluations are scarce (12). Up to now, we only found four publications available (three peer-reviewed), evaluating two assays (15–17, 24), while comparative studies are lacking.

Sensitivity is the most important performance parameter of SARS-CoV-2 antigen detection (12). However, it is also considered the main limiting factor due to experiences with influenza and RSV lateral flow detection kits (6, 25). Newer biosensor-based methods to detect SARS-CoV-2 antigen have shown promising results and could offer a highly sensitive alternative for rapid diagnosis in the future (26, 27).

Our comparison detected significant performance heterogeneity among the evaluated kits. The Bioeasy assay had the highest sensitivity, which was in accordance with our previous evaluation (15). The here reported sensitivity (85%) was almost identical to the value reported by the manufacturer (85.5%). A study from China (with participation of the manufacturer) found an overall sensitivity of 68%; however, for samples with Ct values ≤30, the sensitivity increased to 98% (24). This excellent identification rate in specimens with high viral loads (Ct <25) was confirmed in this report and our recently published study (15). The RapiGen test also showed an acceptable sensitivity (84.9%) with high viral load samples, but was much less sensitive (15.4%) when the viral burden was low. This test had a visual readout, which might have contributed to lower sensitivity. Another assay with visual bands (Respi-Strip CORIS) was recently evaluated in two European studies. Overall sensitivity ranged from 50% to 57.6%; however, detection rates improved for samples with high viral loads (Ct <25) reaching sensitivities of 73.9% to 82.2% (16, 17). The other two assays evaluated by us showed an insufficient performance. Savant had an overall sensitivity of 16.7%, which did not increase significantly in high viral load samples (Figure 2), while the Liming bio assay did not detect any of 10 positives and was not further evaluated.

All four tests had a cassette format; two had visual readout and two were based on immunofluorescence, one of which required an automated reader. In our experience, all systems were easy to use, robust, and gave a qualitative result in short time (10-20 minutes). The fluorescence reader eased interpretation, but it required a higher standard including technical support. The user-friendliness of all tests demonstrated their potential for decentralized screening. However, the application as POCT is hampered by the inherent biological hazard of respiratory specimens, which require processing in a biosafety cabinet (28). This problem could be overcome if extraction buffers or solutions with virus inactivating properties are used.

The enormous performance gaps detected in our study highlight the urgent need of comparative studies of commercialized antigen tests (12). A possible explanation of performance variations might be related to differences in protein targets. However, details on the detection system (target antigen and detecting antibody) are only reported by a minority of manufacturers (29). The methodology of using specimens stored in UTM for evaluation purposes is another critical point with advantages and disadvantages (15), which should be systematically validated, e.g. with spiked samples.

Performance data are critical for local decision making and also for global agencies in the procurement of simpler, scalable diagnostic tests. Although these tests are less sensitive than RT-PCR, they might be useful during outbreak situations, when timely results are important, but access to molecular testing is limited (14). As shown in this and other reports, antigen tests are more reliable in samples with high viral loads, which usually occur during the first days of clinical disease (30–32). In our population, for example, 96% of high viral load specimens derived from patients in their first week of symptoms. Accordingly, the sensitivities for first week samples reached 91%-95% (Bioeasy) in this and previous studies (15). Interestingly, a similar value (94%) was reported for SARS-CoV antigen detection during the first 5 days of disease in a publication from 2004 (33).

The potential to detect early infections might be crucial for the design of new RDT-based algorithms, which are particularly important in weaker health systems and low resource settings such as Latin America and Africa. Possible frontline applications include community-based testing, e.g. in drive-through test stations, and at-home self-testing (34). However, in light of the imperfection of tests, large scale strategies need to be well designed to avoid negative effects (35). Another application might be as an adjunct to RT-PCR to achieve rapid preliminary results, e.g. for healthcare workers.

A limitation regarding the evaluation of specificity was the low number of negative samples and that the study was performed during a season with reduced circulation of other respiratory viruses.

In conclusion, our comparative study highlighted a significant heterogeneity of test performance. The high sensitivity of some tests demonstrated that antigen detection has the potential to serve as an alternative or adjunct diagnostic method to RT-PCR, especially in patients presenting with high viral loads in early phases of infection. This might be of particular importance in situations with limited access to RT-PCR or prolonged turnaround time. Further comparative evaluations are necessary to select for tests with high performance among the growing market of diagnostics for Covid-19.

## Acknowledgements

We thank Savant Biotechnology Co. for providing the test kits free of charge for evaluation.

## Contributors

LP and TW conceived the study. LP, PL, MG, and GP curated the data. LP, TW, PL, and MI analysed the data. LP and GP performed the investigation. LP and TW administered the project. LP, PL, and VV supervised the study. LP, TW, PL, MI, JMM, and RA validated the data. TW wrote the first draft. All authors contributed in reviewing and editing later drafts, and approved the final version.

## References

1. WHO. A coordinated Global Research Roadmap. March 2020. https://www.who.int/blueprint/priority-diseases/key-action/Coronavirus_Roadmap_V9.pdf?ua=1 (accessed 19 May 2020)

2. Nguyen T, Duong Bang D, Wolff A. 2020. 2019 novel coronavirus disease (COVID-19): Paving the road for rapid detection and point-of-care diagnostics. Micromachines (Basel) 11:E306.

3. Corman VM, Landt O, Kaiser M Molenkamp R, Meijer A, Chu DKW, Bleicker T, Brünink S, Schneider J, Schmidt ML, Mulders DG, Haagmans BL, van der Veer B, van den Brink S, Wijsman L, Goderski G, Romette JL, Ellis J, Zambon M, Peiris M, Goossens H, Reusken C, Koopmans MP, Drosten C. 2020. Detection of 2019 novel coronavirus (2019-nCoV) by real-time RT-PCR. Euro Surveill doi:10.2807/1560-7917.ES.2020.25.3.2000045.

4. World Health Organization (WHO). 19 March 2020. Laboratory testing for 2019 novel coronavirus (2019-nCoV) in suspected human cases. Interim guidance. https://www.who.int/publications-detail/laboratory-testing-for-2019-novel-coronavirus-in-suspected-human-cases-20200117 (accessed 19 May 2020)

5. Centers for Disease Control and Prevention, Respiratory Viruses Branch, Division of Viral Diseases. 04 Feb 2020. Real-Time RT-PCR Panel for Detection 2019-Novel Coronavirus. Instructions for Use. https://www.cdc.gov/coronavirus/2019-ncov/lab/index.html?CDC_AA_refVal=https%3A%2F%2Fwww.cdc.gov%2Fcoronavirus%2F2019-ncov%2Flab%2Frt-pcr-detection-instructions.html (accessed 19 May 2020)

6. Cheng MP, Papenburg J, Desjardins M, Kanjilal S, Quach C, Libman M, Dittrich S, Yansouni CP.. 13 April 2020. Diagnostic testing for severe acute respiratory syndrome-related coronavirus-2: A narrative review. Ann Intern Med doi:10.7326/M20-1301

7. World Health Organization (WHO). 22 March 2020. Laboratory testing strategy recommendations for COVID-19. Interim guidance. https://apps.who.int/iris/bitstream/handle/10665/331509/WHO-COVID-19-lab_testing-2020.1-eng.pdf (accessed 19 May 2020)

8. Patel R, Babady E, Theel ES, Storch GA, Pinsky BA, St George K, Smith TC, Bertuzzi S.. 2020. Report from the American Society for Microbiology COVID-19 International Summit, 23 March 2020: Value of diagnostic testing for SARS-CoV-2/COVID-19. mBio 11:e00722–20.

9. Tang YW, Schmitz JE, Persing DH, Stratton CW. 3 April 2020. The laboratory diagnosis of COVID-19 infection: Current issues and challenges. J Clin Microbiol doi: 10.1128/JCM.00512-20.

10. Döhla M, Boesecke C, Schulte B, Diegmann C, Sib E, Richter E, Eschbach-Bludau M, Aldabbagh S, Marx B, Eis-Hübinger AM, Schmithausen RM, Streeck H. 2020. Rapid point-of-care testing for SARS-CoV-2 in a community screening setting shows low sensitivity. Public Health 182:170–172. doi:10.1016/j.puhe.2020.04.009.

11. World Health Organization (WHO). 8 April 2020. Advice on the use of point-of-care immunodiagnostic tests for COVID-19. Scientific Brief. https://www.who.int/news-room/commentaries/detail/advice-on-the-use-of-point-of-care-immunodiagnostic-tests-for-covid-19 (accessed 19 May 2020).

12. European Commission. 15 April 2020. Communication from the Commission: Guidelines on COVID-19 in vitro diagnostic tests and their performance (2020/C 122 I/01). Official Journal of the European Union. https://eur-lex.europa.eu/legal-content/EN/TXT/PDF/?uri=CELEX:52020XC0415(04)&from=EN (accessed 19 May 2020).

13. Foundation for Innovative New Diagnostics (FIND). SARS-CoV-2 diagnostic pipeline. https://www.finddx.org/covid-19/pipeline (accessed 19 May 2020)

14. European Centre for Disease Prevention and Control (ECDC). 23 April 2020. Coronavirus disease 2019 (COVID-19) in the EU/EEA and the UK – ninth update. https://www.ecdc.europa.eu/en/publications-data/rapid-risk-assessment-coronavirus-disease-2019-covid-19-pandemic-ninth-update (accessed 19 May 2020).

15. Porte L, Legarraga P, Vollrath V, Aguilera X, Munita JM, Araos R, Pizarro G, Vial P, Iruretagoyena M, Dittrich S, Weitzel T. 2020. Evaluation of novel antigen-based rapid detection test for the diagnosis of SARS-CoV-2 in respiratory samples. Int J Infect Dis (accepted for publication). http://dx.doi.org/10.2139/ssrn.3569871 (preprint).

16. Mertens P, De Vos N, Martiny D, Jassoy C, Mirazimi A, Cuypers L, Van den Wijngaert S, Monteil V, Melin P, Stoffels K, Yin N, Mileto D, Delaunoy S, Magein H, Lagrou K, Bouzet J, Serrano G, Wautier M, Leclipteux T, Van Ranst M, Vandenberg O and LHUB-ULB SARS-CoV-2 working diagnostic group. 2020. Development and potential usefulness of the COVID-19 Ag Respi-Strip diagnostic assay in a pandemic context. Front. Med 7:225. doi: 10.3389/fmed.2020.00225.

17. Lambert-Niclot S, Cuffel A, Le Pape S, Vauloup-Fellous C, Morand-Joubert L, Roque-Afonso AM, Le Goff J, Delaugerre C; AP-HP/Universities/Inserm COVID-19 research collaboration. 13 May 2020. Evaluation of a rapid diagnostic assay for detection of SARS CoV-2 antigen in nasopharyngeal swab. J Clin Microbiol doi:10.1128/JCM.00977-20.

18. European Centre for Disease Prevention and Control (ECDC). 1 April 2020. An overview of the rapid test situation for COVID-19 diagnosis in the EU/EEA. https://www.ecdc.europa.eu/en/publications-data/overview-rapid-test-situation-covid-19-diagnosiseueea#no-link (accessed 19 May 2020).

19. World Health Organization (WHO). 5 April 2020. Medical Product Alert N°3/2020: Falsified medical products, including in vitro diagnostics, that claim to prevent, detect, treat or cure COVID-19. https://www.who.int/news-room/detail/31-03-2020-medical-product-alert-n-3-2020.

20. Theel ES, Slev P, Wheeler S, Couturier MR, Wong SJ, Kadkhoda K. 29 April 2020. The role of antibody testing for SARS-CoV-2: Is there one? J Clin Microbiol doi:10.1128/JCM.00797-20.

21. Lee CY, Lin RTP, Renia L, Ng LFP. 2020. Serological approaches for COVID-19: Epidemiologic perspective on surveillance and control. Front Immunol 11:879.

22. Whitman JD, Hiatt J, Mowery CT, Shy BR, Yu R, Yamamoto TN, Rathore U, Goldgof GM, Whitty C, Jonathan M. Woo, Gallman AE, Miller TE, Levine AG, Nguyen DN, Bapat SP, Balcerek J, Bylsma S, Lyons AM, Li S, Wong AW, Gillis-Buck EM, Steinhart ZB, Lee Y, Apathy R, Lipke MJ, Smith JA, Zheng T, Boothby IC, Isaza E, Chan J, Acenas II DD, Lee J, Macrae TA, Kyaw TS, Wu D, Ng DL, Gu W, York VA, Eskandarian HA, Callaway PC, Warrier L, Moreno ME, Levan J, Torres L, Farrington L, Loudermilk R, Koshal K, Zorn KC, Garcia-Beltran WF, Yang D, Astudillo MG, Bernstein BE, Gelfand JA, Ryan ET, Charles RC, Iafrate AJ, Lennerz JK, Miller S, Chiu CY, Stramer SL, Wilson MR, Manglik A, Ye CJ, Krogan NJ, Anderson MS, Cyster JG, Ernst JD, Wu AHB, Lynch KL, Bern C, Hsu PD, Marson A. 2020. Test performance evaluation of SARS-CoV-2 serological assays. https://www.medrxiv.org/content/10.1101/2020.04.25.20074856v2 (preprint).

23. Deeks JJ, Dinnes J, Takwoingi Y, Davenport C, Leeflang MMG, Spijker R, Hooft L, Van den Bruel A, Emperador D, Dittrich S. 2020. Diagnosis of SARS-CoV-2 infection and COVID-19: accuracy of signs and symptoms; molecular, antigen, and antibody tests; and routine laboratory markers (Protocol). Cochrane Database of Systematic Reviews doi:10.1002/14651858.CD013596.

24. Diao B, Wen K, Chen J, Liu Y, Yuan Z, Han C, Chen J, Pan Y, Chen L, Dan Y, Wang J, Chen Y, Deng G, Zhou H, Wu Y. Diagnosis of acute respiratory syndrome coronavirus 2 infection by detection of nucleocapsid protein. https://www.medrxiv.org/content/10.1101/2020.03.07.20032524v2 (preprint).

25. Venter M, Richter K. 13 May 2020. Towards effective diagnostic assays for COVID-19: a review. J Clin Pathol doi:10.1136/jclinpath-2020-206685.

26. Seo G, Lee G, Kim MJ, Baek SH, Choi M, Ku KB, Lee CS, Jun S, Park D, Kim HG, Kim SJ, Lee JO, Kim BT, Park EC, Kim SI. 2020. Rapid detection of COVID-19 causative virus (SARS-CoV-2) in human nasopharyngeal swab specimens using field-effect transistor-based biosensor. ACS Nano 14:5135–5142.

27. Mahari S, Roberts A, Shahdeo D, Gandhi S.2020. eCovSens-Ultrasensitive Novel In-House Built Printed Circuit Board Based Electrochemical Device for rapid detection of nCovid-19 antigen, a spike protein domain 1 of SARS-CoV-2. https://www.biorxiv.org/content/10.1101/2020.04.24.059204v3 (preprint).

28. World Health Organization (WHO). 12 February 2020. Laboratory biosafety guidance related to the novel coronavirus (2019-nCoV). Interim guidance. https://www.who.int/docs/default-source/coronaviruse/laboratory-biosafety-novel-coronavirus-version-1-1.pdf?sfvrsn=912a9847_2 (accessed 19 May 2020).

29. González JM, Shelton WJ, Díaz-Vallejo M, Rodriguez-Castellanos VE, Zuluaga JDH, Chamorro DF, Arroyo-Ariza D. 2020. Immunological assays for SARS-CoV-2: an analysis of available commercial tests to measure antigen and antibodies https://doi.org/10.1101/2020.04.10.20061150 (preprint).

30. Wölfel, R., Corman, V.M., Guggemos, W. Seilmaier M, Zange S, Mueller MA, Niemeyer D, Jones TC, Vollmar P, Rothe C, Hoelscher M, Bleicker T, Brünink S, Schneider J, Ehmann R, Zwirglmaier K, Drosten C, Wendtner C. 2020. Virological assessment of hospitalized patients with COVID-2019. Nature https://doi.org/10.1038/s41586-020-2196-x.

31. Zou L, Ruan F, Huang M, Liang L, Huang H, Hong Z, Yu J, Kang M, Song Y, Xia J, Guo Q, Song T, He J, Yen HL, Peiris M, Wu J. 2020. SARS-CoV-2 viral load in upper respiratory specimens of infected patients. N Engl J Med 382: 1177–1179.

32. To KK, Tsang OT, Leung WS, Tam AR, Wu TC, Lung DC, Yip CC, Cai JP, Chan JM, Chik TS, Lau DP, Choi CY, Chen LL, Chan WM, Chan KH, Ip JD, Ng AC, Poon RW, Luo CT, Cheng VC, Chan JF, Hung IF, Chen Z, Chen H, Yuen KY. 2020. Temporal profiles of viral load in posterior oropharyngeal saliva samples and serum antibody responses during infection by SARS-CoV-2: an observational cohort study. Lancet Infect Dis 20:565–574. doi:10.1016/S1473-3099(20)30196-1.

33. Che XY, Hao W, Wang Y, Di B, Yin K, Xu YC, Feng CS, Wan ZY, Cheng VC, Yuen KY. 2004. Nucleocapsid protein as early diagnostic marker for SARS. Emerg Infect Dis 10:1947–1949.

34. Gharizadeh B, Yue J, Yu M, Liu Y, Zhou M, Lu D, Zhang J. 29 April 2020. Navigating the pandemic response life cycle: molecular diagnostics and immunoassays in the context of COVID-19 management IEEE Rev Biomed Eng doi:10.1109/RBME.2020.2991444.

35. Gray N, Calleja D, Wimbush A, Miralles-Dolz E, Gray A, De-Angelis M, Derrer-Merk E, Uchenna Oparaji B, Stepanov V, Clearkin L, Ferson S. 2020. “No test is better than a bad test”: Impact of diagnostic uncertainty in mass testing on the spread of COVID-19. https://www.medrxiv.org/content/10.1101/2020.04.16.20067884v2 (preprint).

